# THETA-Rhythm makes the world go round: Dissociative effects of TMS theta vs alpha entrainment of right pTPJ on embodied perspective transformations

**DOI:** 10.1101/083733

**Authors:** Gerard Gooding-Williams, Hongfang Wang, Klaus Kessler

## Introduction

Being able to imagine another person’s experience and perspective of the world is a crucial human social ability (Call & Tomasello, 1999) and recent reports suggest that humans “embody” another’s viewpoint by mentally rotating their own body representation into the other’s orientation (Kessler, Cao, O’Shea, & Wang, 2014; Kessler & Rutherford, 2010; Kessler & Thomson, 2010; Surtees, Apperly, & Samson, 2013). Using Magnetoencephalography (MEG) our group recently reported in *Cortex* that brain oscillations at theta frequency (3-7 Hz), originating from the right posterior temporo-parietal-junction (pTPJ) reflected cognitive as well as embodied processing elements of perspective transformations (Wang, Callaghan, Gooding-Williams, McAllister, & Kessler, 2016, Expt 1). This was subsequently confirmed using transcranial magnetic stimulation (TMS; Wang et al, 2016, Expt 2) of right pTPJ with a dual pulse protocol (dpTMS), which affected embodied aspects of perspective taking (conforming to other stimulation reports, e.g., Blanke et al., 2005; Santiesteban, Banissy, Catmur, & Bird, 2012; van Elk, Duizer, Sligte, & van Schie, 2016).

Overall these findings emphasised theta oscillations originating from right pTPJ as the core of a wide-spread cortical network involved in embodying another’s viewpoint. However, while dpTMS was suitable for revealing the involvement of right pTPJ, it was insufficient for corroborating the importance of theta oscillations. We therefore employed frequency-specific TMS entrainment (Thut & Miniussi, 2009) in the current study to specifically test the hypothesis that theta entrainment of right pTPJ supports VPT, resulting in faster response times (RTs), while alpha entrainment might inhibit and slow down VPT. The latter was based on the general notion that increased alpha amplitudes reflect inhibition (Jensen & Mazaheri, 2010; Klimesch, Sauseng, & Hanslmayr, 2007) as well as on our observation in Wang et al. (2016) that theta and alpha oscillations fulfil complementary roles, with alpha supporting visual processing at 60°, while theta supporting full-blown VPT at 160°.

## Materials and Methods

### Participants

14 volunteers participated in the experiment (3 left-handed, 8 males, average age 25.0). All recruitment, screening, and experimental procedures complied with the Declaration of Helsinki and were approved by Aston University ethics committee.

### Experimental Procedures

The stimuli and experimental procedures were identical to the dpTMS experiment in Wang et al (2016), with the only difference that instead of 2 TMS pulses administered to right pTPJ during presentation of the perspective taking stimulus (Fig. 1), 15 pulses at either theta frequency (6Hz) or alpha frequency (10Hz) were administered before stimulus onset (e.g. Hanslmayr, Matuschek, & Fellner, 2014). Pulse intensity was 90% of the individual motor threshold, as determined by standard protocols (Rossini et al., 1994). We targeted the identical pTPJ site as in Wang et al., (2016; MNI-coordinates: 50, −60, 32) using Brainsight® neuronavigation and individual MRIs. In addition we followed the rationale of Wang et al. (2016) and employed sham trials (no TMS pulses), where only acoustic clicks were administered via earphones. Click trains were played on all trials at a frequency of 30Hz, thus, participants were unable to distinguish acoustically between 6Hz and 10Hz stimulation and between TMS and sham trials.

**Figure 1.**
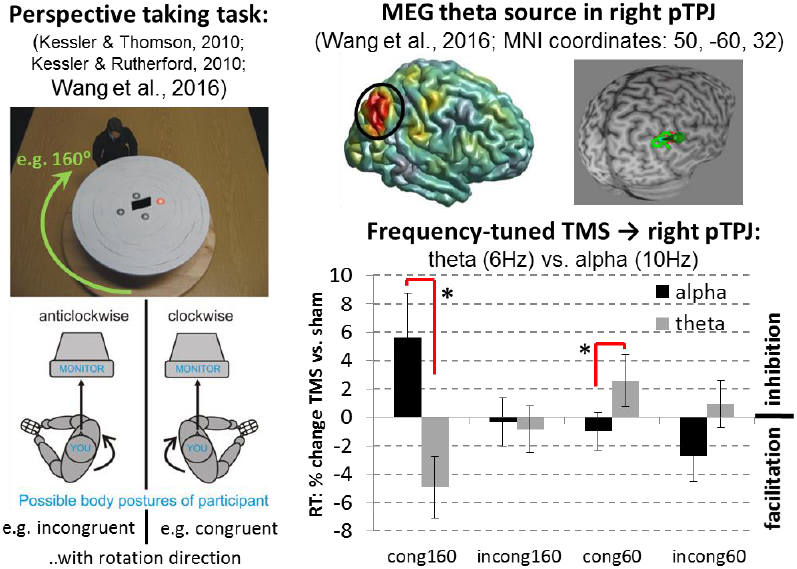
Experimental paradigm adopted from Wang et al. (2016; also Kessler & Rutherford, 2010; Kessler & Thomson, 2010), showing an avatar at a round table sitting at either 60° or 160° angular disparity (clock- or anticlockwise). Participants were instructed to press the left mouse button for a “left” target (red sphere) and the right mouse button for a “right” target in concordance with the avatar’s perspective. Below the stimulus the “posture manipulation” is shown, where the participant’s body was either turned clock- or anticlockwise, while the head remained straight, gazing ahead at the screen. Hence, the body-turn could either be congruent or incongruent with the direction of mental (self-)rotation into the other’s perspective (Kessler & Thomson, 2010). The brain image at the top right shows the pTPJ source obtained for theta oscillations in Wang et al. (2016). Using Brainsight® TMS neuronavigation and individual MRIs (top far-right), the current study employed the shown MNI coordinates as a target for 15 TMS pulses at either theta (6Hz) or alpha (10Hz) frequency, administered before the onset of the avatar stimulus. The graph at the bottom right shows response times (RT) as a percent-change for each TMS condition in relation to its sham baseline. Negative values in the graph indicate facilitation due to TMS entrainment compared to sham, while positive values reflect inhibition. Asterisks (*) indicate significant t-statistics at the 5% level (for *congruent-160°:* t(13)=2.4, p=.03; for *congruent-60°:* t(13)=2.28, p<.04). Error bars denote the standard error of mean.

The same experimental conditions as in Wang et al. (2016, Expt. 2; Fig. 1) were employed with stimuli showing an avatar at a round table sitting at either 60° or 160° angular disparity (clock or anticlockwise) and targets (red sphere) appearing either to the left or right from the avatar’s perspective. Participants were required to press the left mouse button for a “left” target and the right mouse button for a “right” target in concordance with the avatar’s perspective. Figure 1 further shows the unique posture manipulation, which led Kessler and Thomson (2010; replicated in Kessler et al., 2014; Kessler & Rutherford, 2010; Surtees et al., 2013) to conclude that VPT is an embodied process: When the participant’s posture was turned towards the target viewpoint (congruent), perspective transformations were faster than when it was turned in the other direction (incongruent). Wang et al., (2016) replicated this behavioural effect in their MEG experiment and in the TMS sham condition, while dpTMS to right pTPJ abolished the posture effect.

Our design included four repeated measures factors (angular disparity: 60°/160°; posture: congruent/incongruent; TMS frequency: theta/alpha; TMS condition: 15pulses/sham) and 20 trials per condition were administered (320 trials in total). Conditions were presented randomly apart from posture and frequency. A particular posture (e.g. turned clockwise) was adopted throughout a mini-block of 16 trials and then changed to the other posture for the next mini-block. A single TMS entrainment frequency, theta or alpha, was applied throughout a block of 32 trials (2 mini-blocks).

## Results and Discussion

We calculated the percent-change for each TMS condition in relation to its sham baseline and then conducted a 3-way repeated measures ANOVA. We observed a significant interaction between frequency and angular disparity (F(1, 13)=7, p=.02, η^2^p=.351) indicating that the two entrainment frequencies had opposite effects (Figure 1). Conforming to our hypothesis, theta facilitated VPT at 160°, while alpha slowed it down, and the reverse was true at 60°. This differential effect was more pronounced for a congruent posture as revealed by post-hoc t-tests (Fig. 1) and a statistical trend for a 3-way interaction frequency x angle x posture (F(1, 13)=3.8 p=.074, η^2^p=.225). The facilitatory effect of alpha at 60° compared to 160° is also noteworthy, suggesting that inhibition of embodied processing in pTPJ (alpha entrainment) at 60° might facilitate a visual processing strategy: At 60° the target is still visibly left or right from an egocentric perspective, thus, a perspective transformation is not strictly necessary. For discussion see Kessler & Thomson, (2010) and Wang et al. (2016).

In conclusion our results further corroborate our previous MEG finding that theta oscillations in the right pTPJ are of significance to embodied perspective transformations. As discussed in detail in Wang et al. (2016; also Kessler & Braithwaite, 2016), we propose that pTPJ not only controls representations of self vs. “other” (e.g., Santiesteban et al., 2012) but might actually control the simulation of an alternative embodied self that serves as the basis for inferring other’s mental representations (e.g. their perspective).

## References

Blanke, O., Mohr, C., Michel, C. M., Pascual-Leone, A., Brugger, P., Seeck, M., … Thut, G. (2005). Linking out-of-body experience and self processing to mental own-body imagery at the temporoparietal junction. J Neurosci, 25(3), 550–557.

Call, J., & Tomasello, M. (1999). A nonverbal false belief task: The performance of children and great apes. Child Development, 70(2), 381–395.

Hanslmayr, S., Matuschek, J., & Fellner, M.-C. (2014). Entrainment of prefrontal beta oscillations induces an endogenous echo and impairs memory formation. Current Biology, 24(8), 904–909.

Jensen, O., & Mazaheri, A. (2010). Shaping functional architecture by oscillatory alpha activity: gating by inhibition. Frontiers in human neuroscience, 4, 186.

Kessler, K., & Braithwaite, J. J. (2016). Deliberate and spontaneous sensations of disembodiment: capacity or flaw? Cognitive Neuropsychiatry, 1–17.

Kessler, K., Cao, L., O’Shea, K. J., & Wang, H. (2014). A cross-culture, cross-gender comparison of perspective taking mechanisms. Proceedings of the Royal Society of London B: Biological Sciences, 281(1785), 20140388.

Kessler, K., & Rutherford, H. (2010). The two forms of Visuo-Spatial Perspective Taking are differently embodied and subserve different spatial prepositions. [Original Research]. Frontiers in Psychology, 1. doi: 10.3389/fpsyg.2010.00213

Kessler, K., & Thomson, L. A. (2010). The embodied nature of spatial perspective taking: Embodied transformation versus sensorimotor interference. Cognition, 114(1), 72–88.

Klimesch, W., Sauseng, P., & Hanslmayr, S. (2007). EEG alpha oscillations: The inhibition-timing hypothesis. Brain Research Reviews, 53(1), 63–88.

Rossini, P. M., Barker, A., Berardelli, A., Caramia, M., Caruso, G., Cracco, R., … Lücking, C. (1994). Noninvasive electrical and magnetic stimulation of the brain, spinal cord and roots: basic principles and procedures for routine clinical application. Report of an IFCN committee. Electroencephalography and clinical neurophysiology, 91(2), 79–92.

Santiesteban, I., Banissy, M. J., Catmur, C., & Bird, G. (2012). Enhancing social ability by stimulating right temporoparietal junction. Current Biology, 22(23), 2274–2277.

Surtees, A., Apperly, I., & Samson, D. (2013). The use of embodied self-rotation for visual and spatial perspective-taking. Frontiers in human neuroscience, 7.

Thut, G., & Miniussi, C. (2009). New insights into rhythmic brain activity from TMS–EEG studies. Trends in cognitive sciences, 13(4), 182–189.

van Elk, M., Duizer, M., Sligte, I., & van Schie, H. (2016). Transcranial direct current stimulation of the right temporoparietal junction impairs third-person perspective taking? Cognitive, Affective, & Behavioral Neuroscience, 1–15.

Wang, H., Callaghan, E., Gooding-Williams, G., McAllister, C., & Kessler, K. (2016). Rhythm makes the world go round: An MEG-TMS study on the role of right TPJ theta oscillations in embodied perspective taking? Cortex, 75, 68–81.

